# Mapping of m^6^A and Its Regulatory Targets in Prostate Cancer Reveals a METTL3-low Induction of Therapy Resistance

**DOI:** 10.1101/2021.01.12.426354

**Authors:** Kellie A. Cotter, John Gallon, Nadine Uebersax, Philip Rubin, Kate D. Meyer, Salvatore Piscuoglio, Samie R. Jaffrey, Mark A. Rubin

## Abstract

Recent evidence has highlighted the role of *N*^6^-methyladenosine (m^6^A) in the regulation of mRNA expression, stability and translation, supporting a potential role for post-transcriptional regulation mediated by m^6^A in cancer. Here we explore prostate cancer as an exemplar and demonstrate that low levels of *N*^6^-adenosine-methyltransferase (*METTL3*) is associated with advanced metastatic disease. To explore this relationship, we generated the first prostate m^6^A maps, and further examined how *METTL3* regulates expression at the level of transcription, translation, and protein. Significantly, transcripts encoding extracellular matrix proteins are consistently upregulated with *METTL3* knockdown. We also examined the relationship between *METTL3* and androgen signaling and discovered the upregulation of a hepatocyte nuclear factor-driven gene signature that is associated with therapy resistance in prostate cancer. Significantly, *METTL3* knockdown rendered the cells resistant to androgen receptor antagonists, implicating changes in m^6^A as a mechanism for therapy resistance in metastatic prostate cancer.

## BACKGROUND

At the molecular level, prostate cancer (PCa) is generally characterized by large-scale structural genomic rearrangements^1,2^ and relatively few recurrent genomic alterations^3,4^, which thus far are insufficient to explain the heterogeneity of the disease, and suggest the importance of other mechanisms for gene regulation. In particular, multi-omics studies in PCa have demonstrated that for many oncogenes copy number alterations, DNA methylation, and even mRNA abundance are insufficient to explain the variation seen in protein expression^5,6^, emphasizing a sizeable role for post-transcriptional gene regulation in PCa.

Analogous to epigenetic modifications of DNA, mRNA is also subject to multiple biochemical modifications, the most prevalent being *N*^6^-methyladenosine (m^6^A)^7^. m^6^A is a dynamic mRNA modification and is regulated by the methyltransferase “writer” complex (METTL3, METTL14, WTAP, VIRMA, ZC3H13 and RBM15)^8-12^ the demethylase “eraser” (ALKBH5)^13^ and “reader” proteins (YTHDF1/2/3)^14,15^. m^6^A is found on ∼25% of all transcripts, and is unevenly distributed, with the majority of residues localized near the stop codon or in 3’ untranslated regions (UTRs). Mechanistically, m^6^A has been shown to both destabilize mRNA transcripts^9,15,16^ and increase translation^17^, in addition to influencing splicing^14,18^ and miRNA processing^19^.

Alterations in m^6^A have been associated with progression of multiple cancer types. In glioblastoma and breast cancer, increased expression of the demethylase ALKBH5 and concurrent reduced levels of m^6^A were implicated in the promotion of tumorigenesis and self-renewal of stem-like cells by stabilizing key mRNAs (e.g. the pluripotency factor *NANOG*)^20-22^. In contrast, in lung adenocarcinoma and myeloid leukemia, elevated expression of METTL3 promoted the translation efficiency of several important oncogenes, and enhanced cell growth and invasion^23,24^. Lastly, *METTL3* has been shown to be both highly expressed in prostate cancer, and essential for proliferation in multiple prostate cancer cell lines^25-27^, strongly supporting the significance of m^6^A methylation in PCa.

Of particular concern clinically is metastatic, castration-resistant PCa (CRPC) in which the disease develops resistance to both first-line treatment with androgen deprivation therapy, and to second generation androgen receptor (AR) signaling inhibitors (ARSi, ie. abiraterone or enzalutamide)^28^. There are several proposed mechanisms of resistance that lead to CRPC including alterations in AR which permit activity in low-androgen settings, alterations in pathways downstream of AR, and increased signaling through other signaling pathways allowing for AR independence^29^. Despite some recent progress in developing new treatments for CRPC, average survival at this advanced disease stage is approximately 3 years^30^. Determining the mechanisms behind ARSi resistance are essential to improve existing therapies and finding new potential drug targets.

To address this gap in knowledge, we investigated the role of m^6^A and METTL3 in the progression of PCa, with a particular focus on its role in regulating AR signaling and the response to ARSi. We show that low levels of *METTL3* in patients is associated with clinical markers of CRPC, in particular dysregulation of AR signaling. Using miCLIP we mapped which transcripts are marked by m^6^A, and then further examined how METTL3 knockdown alters expression at the level of transcript, translation and protein. In comparing these cell line results to patient data we find that low METTL3 consistently increases expression of extracellular matrix genes. Furthermore, we examined the relationship between METTL3 and AR signaling and show that while knockdown of METTL3 has no effect on AR response genes, it does render the cells resistant to ARSi in an AR-independent manner. Overall, these findings support a new model for m^6^A function in affecting therapeutic sensitivity to ARSi, and suggests that patients with low levels of *METTL3* expression may not demonstrate an optimal response from ARSi.

## RESULTS

### Low *METTL3* expression is associated with advanced prostate cancer

Given the ability of m^6^A to influence gene expression, we asked whether m^6^A affects PCa progression. To determine if changes m^6^A might contribute to prostate cancer in patients, we assessed if the expression of the methyltransferase m^6^A “writer” complex members (*METTL3, METTL14, WTAP, VIRMA, ZC3H13* and *RBM15*) is related to any clinical parameters in advanced PCa. In a recent precision oncology study, we reported on 430 men with castration resistant prostate cancer (CRPC)^31^. We subset these patients based on the relative expression levels of each member into low (z-score of less than −1) and high (z-score of greater than 1) groups. We then assessed whether these groups demonstrated low androgen receptor (AR) signaling or high neuroendocrine scores, both important measures for a lack of response to anti-androgen therapy^32^.

Although previous studies suggested that *METTL3* expression is higher in primary PCa as compared to benign prostate tissue^26,27^, our examination of the expression of *METTL3* in metastatic PCa samples revealed the opposite phenomena, that decreased *METTL3* is associated with the most aggressive subclass of advanced prostate disease. In this cohort of 430 men with metastatic CRPC, we observed that those with lower levels of *METTL3* demonstrated significantly disrupted AR signaling (*P* = 0.0011) (Figure 1A), and significantly increased expression of genes associated with neuroendocrine progression (*P* = 0.0214) (Figure 1B). This association between low expression and low AR score also held true for *METTL14* (*P* = 0.0002), *WTAP* (*P* = 0.0002), and *VIRMA* (*P* = 3.79E-06) (Supplementary Figure 1A). In contrast *ZC3H13* expression did not show a significant with AR score (Supplementary Figure 1A), however it was the only other gene besides *METTL3* to have a significant association with NEPC score, albeit in the opposite direction (*P* = 0.0433) (Supplementary Figure 1B).

**Figure 1.**
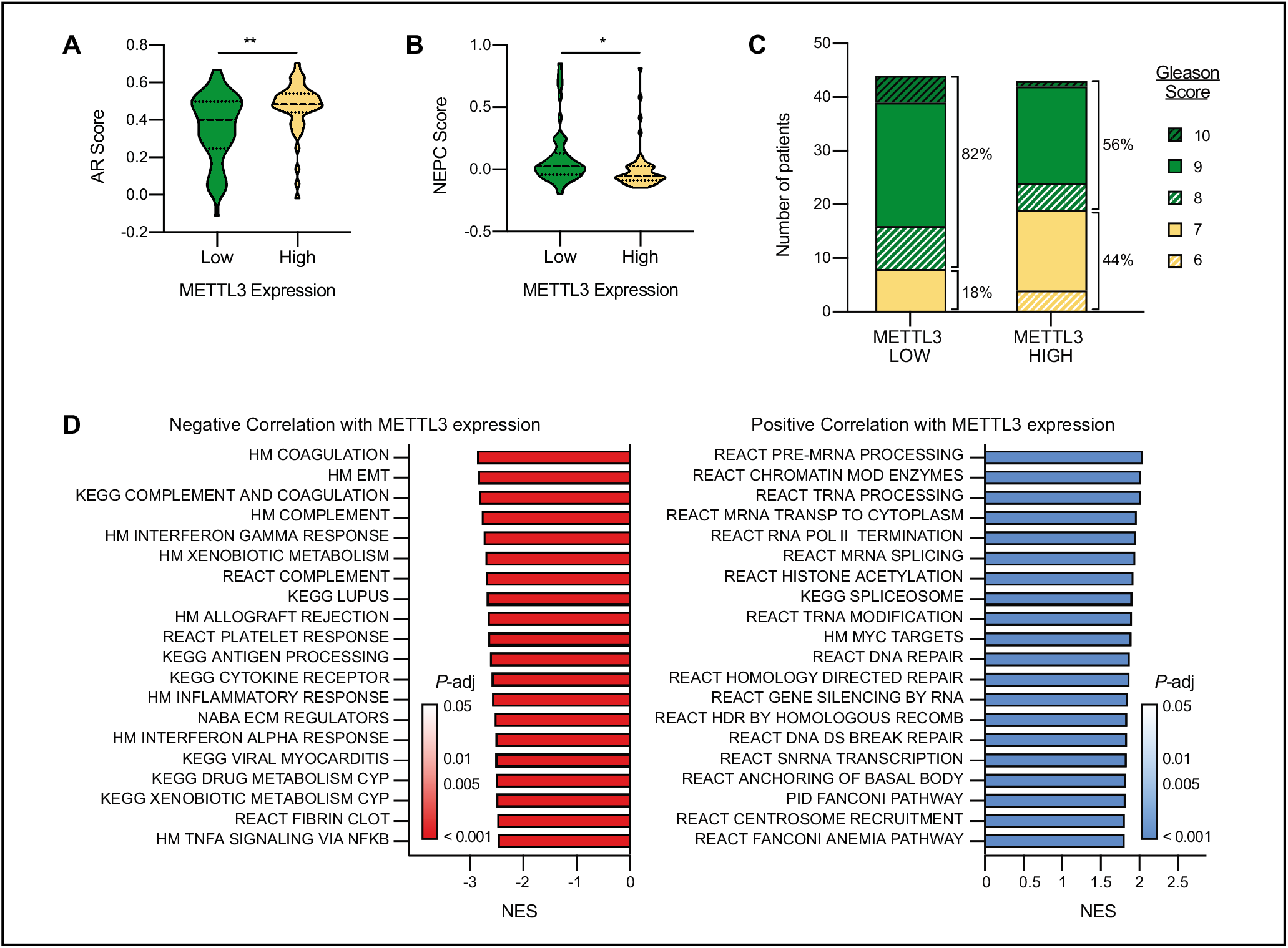
Low *METTL3* expression is associated with advanced-stage disease. Castration resistant prostate cancer samples^31^ were grouped based on *METTL3* expression: high, z-score > 1, n = 50; low, z-score < −1, n = 49. **A)** AR score in *METTL3* high vs. low samples. Median and quartiles are indicated with a dotted line, *P* = 0.0011 as determined by t-test. **B)** NEPC score in *METTL3* high vs. low samples. Median and quartiles are indicated with a dotted line, *P* = 0.0214 as determined by t-test. **C)** Gleason score in *METTL3* high vs. low samples. *P* = 0.0111 as determined by Fischer’s exact test between samples with Gleason 6-7 and Gleason 8+ scores. **D)** Gene set enrichment analysis of genes differentially expressed in in *METTL3* high vs. low samples. Genes were ranked according to the log of the Benjamini and Hochberg adjusted *P*-value as determined by t-test. Shown are the top 20 results in either direction.

In this same group of patients those with lower *METTL3* expression were also more likely to have primary tumors with higher Gleason scores (8+) than those with high *METTL3* expression (*P* = 0.0111) (Figure 1C). None of the other methyltransferase complex members showed any association with Gleason score (Supplementary Figure 1C). Given these data, and its known role as the catalytic subunit of the “writer” complex we chose to focus further investigations on *METTL3*.

To begin to understand the mechanistic links between m^6^A and these signifiers of aggressive CRPC, we asked what genes and/or pathways are regulated by METTL3 in PCa. To begin to address this question we took the same cohort of metastatic patient samples categorized into *METTL3*-high and *METTL3*-low groups, and looked for genes over- or under-expressed in one group versus the other. This analysis identified 2692 genes with significantly reduced expression in the *METTL3*-low group and 2081 genes with significantly increased expression. Gene set enrichment analysis of these genes demonstrates that those that correlate positively with *METTL3* expression mainly involve genes associated with transcription and mRNA processing (Figure 1D). On the other hand, gene sets that display a negative correlation with *METTL3* expression include many pathways implicated in aggressive metastatic disease, including complement and coagulation pathways, EMT, regulation of the extracellular matrix, and drug metabolism (Figure 1D).

### Mapping m^6^A in prostate cancer

Given that our above analysis relies solely on correlations in gene expression within a diverse group of patient samples, we aimed to more thoroughly define the role of m^6^A methylation in the regulation of PCa gene expression. As a first step we sought to generate the first epitranscriptomic maps of m^6^A in PCa. Using m^6^A individual-nucleotide-resolution cross-linking and immunoprecipitation (miCLIP)^33^ we mapped the location of m^6^A at a single-nucleotide resolution throughout the entire transcriptomes of both the AR-sensitive adenocarcinoma LNCaP and in the benign RWPE cell line and identified over 18000 methylated residues in 6653 transcripts. Similar to previously reported m^6^A maps generated in other cell lines and tissues the majority of m^6^A sites identified in both cell lines were in the 3’ UTR, particularly in the vicinity of the stop codon (Figure 2A). Furthermore, the sequence context surrounding the methylation sites was also consistent with the previously reported DRACH motifs (D= A/G/U, R= A/G, H= A/C/U)^14,34^ (Figure 2B).

**Figure 2.**
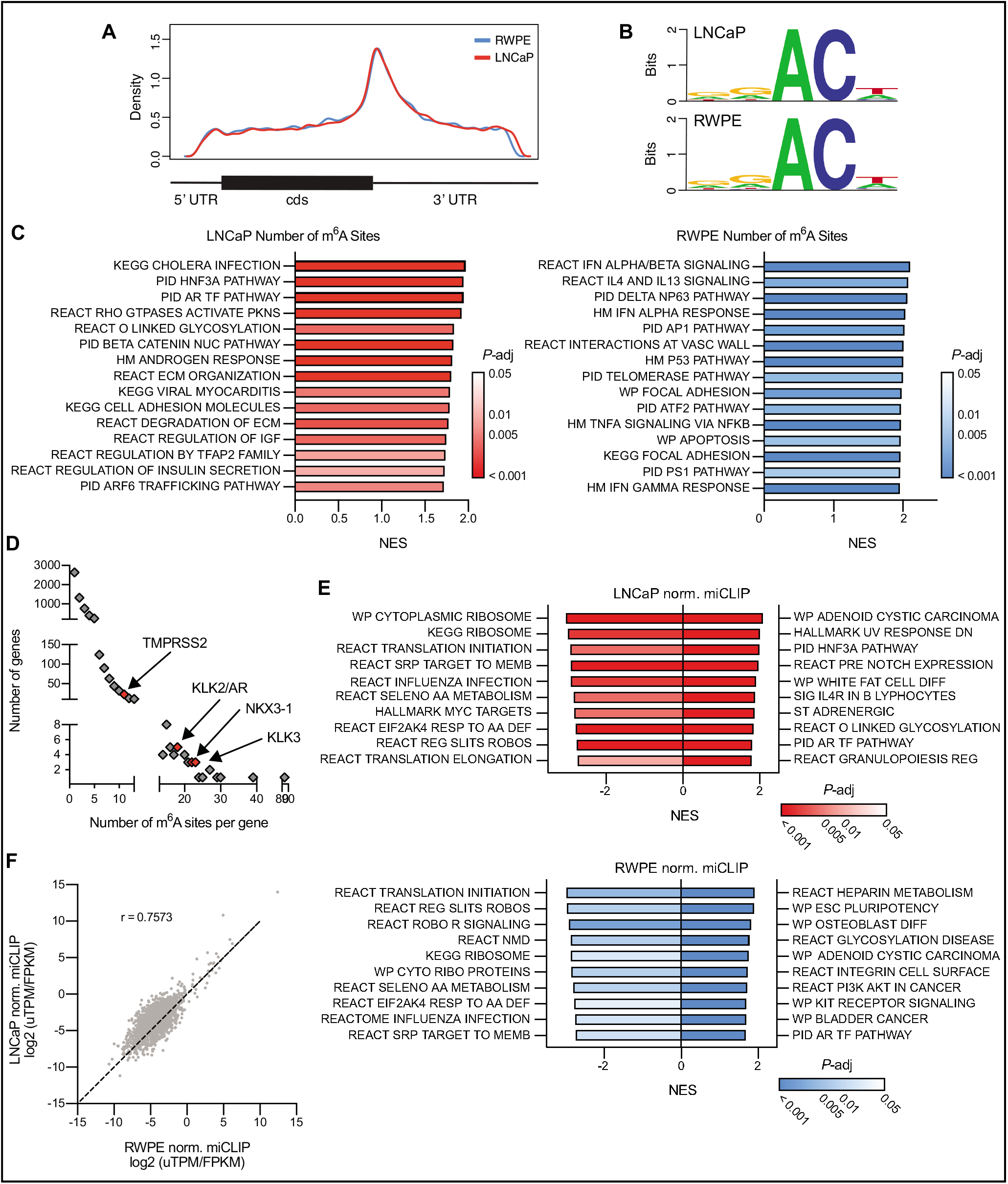
Mapping genes methylated with m^6^A in prostate cancer. m^6^A was mapped in LNCaP and RWPE cell lines using miCLIP. **A)** Metagene plot of the distribution of m^6^A residues along the transcript in both cell lines. **B)** Consensus sequence surrounding m^6^A sites identified in both cell lines generated with WebLogo. **C)** Gene set enrichment analysis of genes ranked by the number of m^6^A sites. Shown are the top 15 gene sets. **D)** AR signature genes (n = 30; Hieronymus, 2006) are significantly (*P* = 0.0004) enriched in m^6^A sites (>10 sites) as determined by Fisher’s exact test. **E)** Gene set enrichment analysis of genes ranked by the total normalized miCLIP unique tags per million for a given transcript. **F)** Comparing the normalized miCLIP unique tags per million per transcript in genes which are similarly expressed (< 10-fold difference) between the LNCaP and RWPE call lines.

In both lines the 70-85% of transcripts had only one or two sites, however 1-2% of the transcripts had 10 or more sites (Supplementary Data 1, 2). In both lines the non-coding RNA *MALAT1* was the most highly methylated with 84 sites in LNCaP and 69 in RWPE. Examination of the association between the total of number of sites identified per transcript and its expression shows that while there is some bias towards highly expressed genes having more total sites (due to inherent biases in IP-based protocols), expression is not the sole driver of the differences seen (Supplementary Figure 2A). Furthermore, there was no correlation between the number of m^6^A sites and the transcript length (Supplementary Figure 2B).

To determine the cellular pathways that m^6^A might influence in prostate cancer, we interrogated the miCLIP data using gene set enrichment analysis to identify pathways enriched in the transcripts that exhibited higher number (i.e., greater than 5) of m^6^A sites. This analysis implicated m^6^A in regulating mRNAs that encode pathways related to cell adhesion and the extracellular matrix in both cell lines (Figure 2C). Remarkably, this enrichment is similar to the gene expression changes linked to low *METTL3* expression based on examination of the patient samples (see Figure 1D). Additionally, this analysis identified AR-regulated genes as also having a high number of m^6^A sites in LNCaP cells. In particular AR signature genes including *KLK2, KLK3, NKX3-1* and even *AR* itself all had greater than 10 sites (Figure 2D).

We further characterized the methylation status of transcripts by not only the number of sites on a given transcript, but also by a relative measurement of the amount of m^6^A at a given site. To do so we normalized the amount fragmented RNA immunoprecipitated using the m^6^A antibody from each individual site to the library size (uTPM), and overall expression of that transcript (Supplementary Figure 2C). As in Supplementary Figure 2A, we see a correlation between the amount of immunoprecipitated RNA and the expression of the transcript. Nonetheless we can also identify distinct subsets of transcripts which have relatively high or low levels of methylation given their expression. In both cell lines this analysis again identified AR pathway genes as having high levels of methylation, whereas ribosomal and translation pathways had low levels of methylation relative to the expression levels of those transcripts (Figure 2E) in agreement with results from other human cell lines^11^. Comparisons of methylation sites between the two lines is of course heavily biased by the highly divergent transcriptomes between LNCaP and RWPE cells. However, upon limiting the analysis to transcripts with less than a 10-fold difference in expression between the two lines we find a strong correlation between the amount of m^6^A (Pearson r = 0.7573) (Figure 2F). In particular, upon comparing the amount of methylation at m^6^A sites common to both cell lines we found a striking concordance between the levels in LNCaP and RWPE cells (Pearson r = 0.8666) (Supplementary Figure 2D).

### METTL3 regulation of gene expression and translation

Although the miCLIP analysis identified many highly methylated transcripts with known functional relevance in PCa, in particular *AR* and its downstream target genes, it remains to be seen whether methylation of these transcripts plays a specific role in their regulation, and further whether this regulation is at the level of transcript stability and/or translation efficiency. To address this, we generated LNCaP cell lines infected with drug-inducible shRNAs targeting *METTL3*. With these lines we were able to precisely and coordinately manipulate both the timing and extent of reduction of *METTL3* expression. We demonstrated a robust knockdown of *METTL3* expression after 96 hours of doxycycline treatment (Figure 3A) as compared to a control non-targeting shRNA against GFP. Further the *METTL3* knockdown is reversible after 144 hours of doxycycline wash-out (Supplementary Figure 3A).

**Figure 3.**
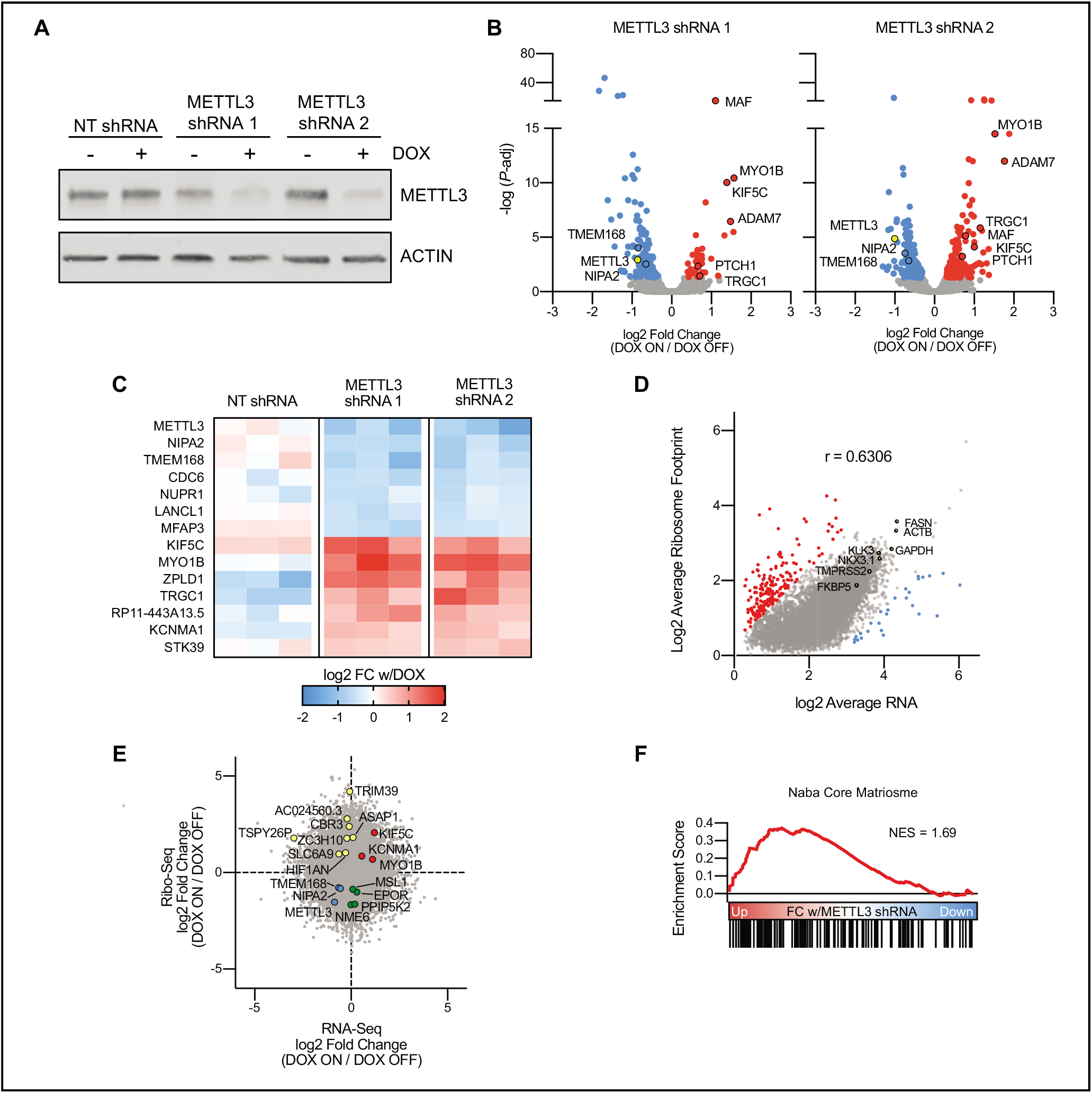
Knockdown of *METTL3* in prostate cancer alters gene expression and translation. **A)** Generation of LNCaP cell lines with two doxycycline-inducible shRNAs targeting *METTL3*, or a non-targeting shRNA targeting *GFP*. Cells were treated with doxycycline for 96 hours followed by Western blot analysis for METTL3. **B)** Significantly (adjusted *P*-value < 0.05) differentially expressed genes with *METTL3* knockdown in the two inducible shRNA lines (n = 3). Common genes between the two shRNA lines are highlighted with a black border. **C)** Six transcripts are significantly down-regulated and seven transcripts are significantly up-regulated with *METTL3* knockdown in both shRNA lines, but not the non-targeting line (n = 3). **D)** Correlation (Pearson r = 0.6306, *P* < 1E-15) between the average mRNA (n = 6) and ribosome footprint (n = 4) from the *METTL3* shRNA lines without doxycycline. Outliers with extreme high (red) and low (blue) translation efficiency (Ribo/RNA) were calculated using ROUT analysis (Q = 1%). **E)** Riborex analysis of Ribo-seq data identities genes with changes in TE independent of changes in mRNA expression. Shown is the fold change with doxycycline treatment in both *METTL3* shRNAs combined as determined by DESeq2 (RNA-seq n = 12, Ribo-Seq n = 10). **F)** Gene set enrichment analysis of genes ranked by the fold change in TE with *METTL3* knockdown as determined by Riborex.

Using these inducible lines, we measured changes in both expression and translation efficiency by means of tandem RNA-seq and ribosome footprint profiling (Ribo-Seq). RNA-seq analysis demonstrated relatively modest changes in gene expression with *METTL3* knockdown. With either *METTL3* shRNA we found 187 and 337 significantly (FDR < 0.05) differentially expressed genes between the doxycycline on and off conditions (Figure 3B). However, we also identified 508 significant gene expression changes in the non-targeting line in response to doxycycline, including three which were also significant in both our *METTL3* shRNA lines (Supplementary Figure 3B). This strategy allowed us to specifically identify six down- and seven up-regulated transcripts (Figure 3C) which were significantly regulated in two different *METTL3* knockdown lines, after considering non-specific effects of doxycycline treatment or shRNA induction.

To determine if these changes persisted at the protein level, we validated one of the most up-regulated transcripts *KIF5C*. We confirmed persistent KIF5C upregulation at the protein level after only 72 hours of doxycycline treatment (Supplementary Figure 3C). *KIF5C* is further of interest in that in metastatic PCa samples its expression is highly correlated with androgen receptor signaling (Spearman’s ρ = 0.56) (Supplementary Figure 3D). Given that expression of *KIF5C* itself is not regulated by androgen receptor signaling in LNCaP cells (Fold-change < 2 with DHT or ENZ) (Supplementary Figure 3E), it remains to be seen whether KIF5C operates upstream of AR, or if it plays some other role in PCa progression.

Similar to published Ribo-seq results from other cell lines and tissues^35^ ribosome footprinting read counts generally correlated with mRNA expression in LNCaP cells (Pearson r = 0.6306) (Figure 3D). Among the most highly expressed and ribosome-bound transcripts are the housekeeping genes *GAPDH* and *ACTB*, and the known AR-targets *KLK3, FASN, NKX3*.*1, FKBP5*, and *TMPRSS2*. We did, however, also identify 225 transcripts that were outliers (ROUT analysis, Q = 1%) with ribosome footprints that do not correlate with mRNA expression. 8 of the 27 transcripts with exceptionally low translation efficiency (TE) are mitochondrial genes which are known to be inefficiently captured with standard ribosome footprinting protocols^36^. Other gene sets with generally low TE include ribosomal proteins and pathways involved in translation (similar to results from proteogenomic studies demonstrating a low correlation between transcript and protein for ribosomal proteins in human tumors^37-39^), while those with high TE are predominated by extracellular matrix proteins in particular collagens (Supplementary Figure 3F).

We then utilized Riborex^40^ analysis to identify genes with significant changes in TE independent of any changes in mRNA expression. This analysis further identified 15 and 6 genes with a significant (FDR < 0.1) increase or decrease in TE with *METTL3* knockdown in both of the inducible shRNA lines (Figure 3E). Gene set enrichment analysis demonstrated that the TE of transcripts encoding extracellular matrix proteins is increased with *METTL3* knockdown (Figure 3F).

### METTL3 regulation of protein expression

Given the changes we detected in translation efficiency with *METTL3* knockdown we then wanted to understand how these changes are reflected in protein abundance. Therefore, we conducted shot-gun proteomics using the same inducible *METTL3* shRNA lines in order to get an unbiased and quantitative picture of the entire proteome. Protein expression generally correlated with mRNA expression (Spearman’s ρ = 0.443-0.469) in line with recent results describing paired RNA-seq and proteomics in PCa patient samples^6^ (Figure 4A).

**Figure 4.**
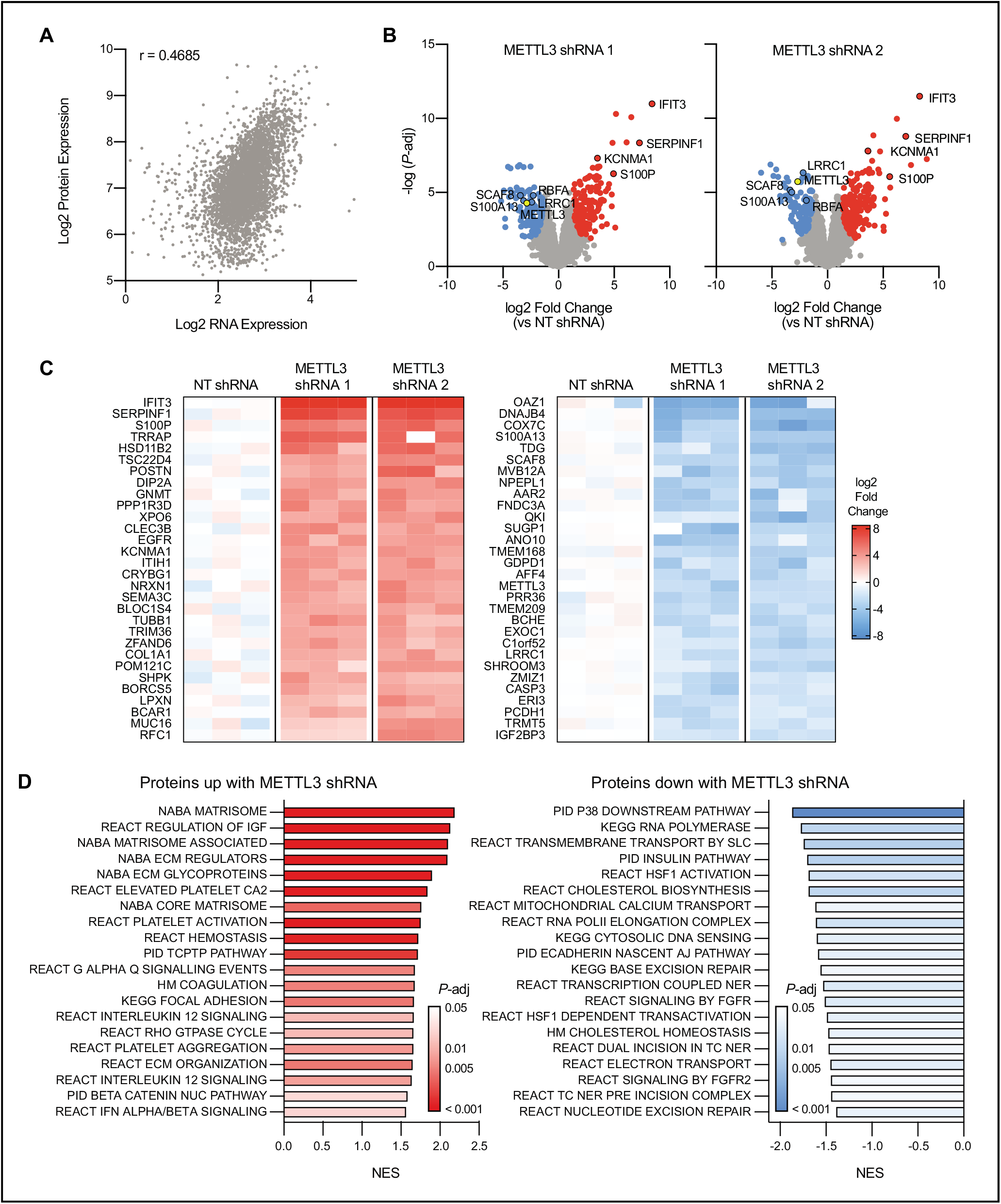
Knockdown of *METTL3* in prostate cancer alters protein expression. **A)** Correlation (Spearman’s ρ = 0.4685, *P* < 1E-15) between the average mRNA expression (n = 3) and protein expression (n = 3) as determined by shotgun proteomics in the NT shRNA line. **B)** Significantly (adjusted *P*-value < 0.05) differentially expressed proteins with *METTL3* knockdown in the two inducible shRNA lines as compared to the NT shRNA line (n = 3 for each shRNA). Common highly differentially expressed proteins between the two shRNA lines are highlighted with a black border. **C)** The top 60 differentially expressed proteins seen with *METTL3* knockdown in both shRNA lines as compared to the NT control (n = 3 for each shRNA). **D)** Gene set enrichment analysis of proteins differentially expressed with *METTL3* knockdown. Proteins were ranked according to the log of the adjusted *P*-value as determined by t-test. Shown are the top 20 results in either direction.

We identified 527 and 537 significantly differentially expressed (FDR adj p-value < 0.05, FC > 2) proteins in either *METTL3* shRNA line as compared to the non-targeting shRNA line (Figure 4B). Similar to our Ribo-seq results we also observed much larger fold changes at the protein level (> 15-fold) than we had seen at the level of expression (2-4-fold, Figure 3B). Unfortunately, only one of the transcripts with significant changes in TE were detected in our proteomics experiment, thus precluding a thorough comparison between the two methods; nonetheless this protein (CBR3) was upregulated in both cell lines after *METTL3* knockdown. In total 182 proteins significantly upregulated in both *METTL3* shRNA lines while 109 proteins were significantly down-regulated (Figure 4C, Supplementary Data 3). Previous studies have identified interferon signaling as being highly upregulated in response to *METTL3* knockdown^41,42^. Here we identified an upregulation of the interferon-induced proteins IFIT1, IFIT2, and IFIT3, but not other interferon-induced proteins. This is likely due to the fact that LNCaP cells are unresponsive to interferon signaling (Supplementary Figure 4) due to a silencing of JAK1 by bi-allelic inactivating mutations and epigenetic silencing in this line^43,44^.

Gene set enrichment analysis of the proteomics data revealed that down-regulated proteins mainly included those associated with transcription and repair of DNA damage (Figure 4D). Gene sets with up-regulated proteins predominantly consisted of those involved in the composition and regulation of the extracellular matrix, coagulation and cell adhesion. This is consistent with results seen in patients with low *METTL3* expression (Figure 1D) and genes with high levels of m^6^A in LNCaP cells (Figure 2D).

### *METTL3* knockdown triggers AR-independence

Recent work has demonstrated that the majority of the effects of m^6^A is mediated through a reduction in transcript stability^16,45^. Given this, one would expect that we would see an increase in expression of the highly methylated AR pathways genes (Figure 2D), and therefore AR signaling, with *METTL3* knockdown. In contrast, we see no change in the expression of these genes at the transcript (Figure 3C) or protein level (Figure 4C). Furthermore, the expectation of an increase in AR signaling with *METTL3* knockdown is inconsistent with the results we saw in the metastatic patient cohort where low levels of *METTL3* were associated with a lower AR score (Figure 1A).

To resolve these contradictions, we decided to directly examine the response of the *METTL3* knockdown lines to treatment with enzalutamide (ENZ, a potent AR ligand binding inhibitor), and/or stimulation with dihydroxytestosterone (DHT). We then examined the changes in expression of known AR target genes (i.e., *KLK3, TMPRSS2, FKBP5* and *NKX3*.1) by qPCR (Figure 5A). In the absence of *METTL3* knockdown all three cell lines responded similarly to the treatments. Further, with *METTL3* knockdown we saw no change in the response of *KLK3, FKBP5* and *TMPRSS2* to AR inhibition with ENZ or charcoal-stripped (hormone-free) media, nor to DHT stimulation. In contrast, we did see a significant reduction in the down-regulation of *NKX3*.*1* with *METTL3* knockdown and ENZ (sh1, *P* = 0.0062; sh2, *P* = 0.0074) or charcoal-stripped media (sh1, *P* = 0.036; sh2, *P* = 0.044) treatment. *NKX3*.*1* is a prostate-specific homeobox gene that regulates normal prostate differentiation and suppresses PCa initiation^46^. The upregulation of *NKX3*.*1* with DHT stimulation, however, remained unchanged (FC sh1 = 2.3, FC sh2 = 2.6, vs FC NT = 3). As such, we concluded that *METTL3* knockdown does not alter AR signaling overall in LNCaP cells.

**Figure 5.**
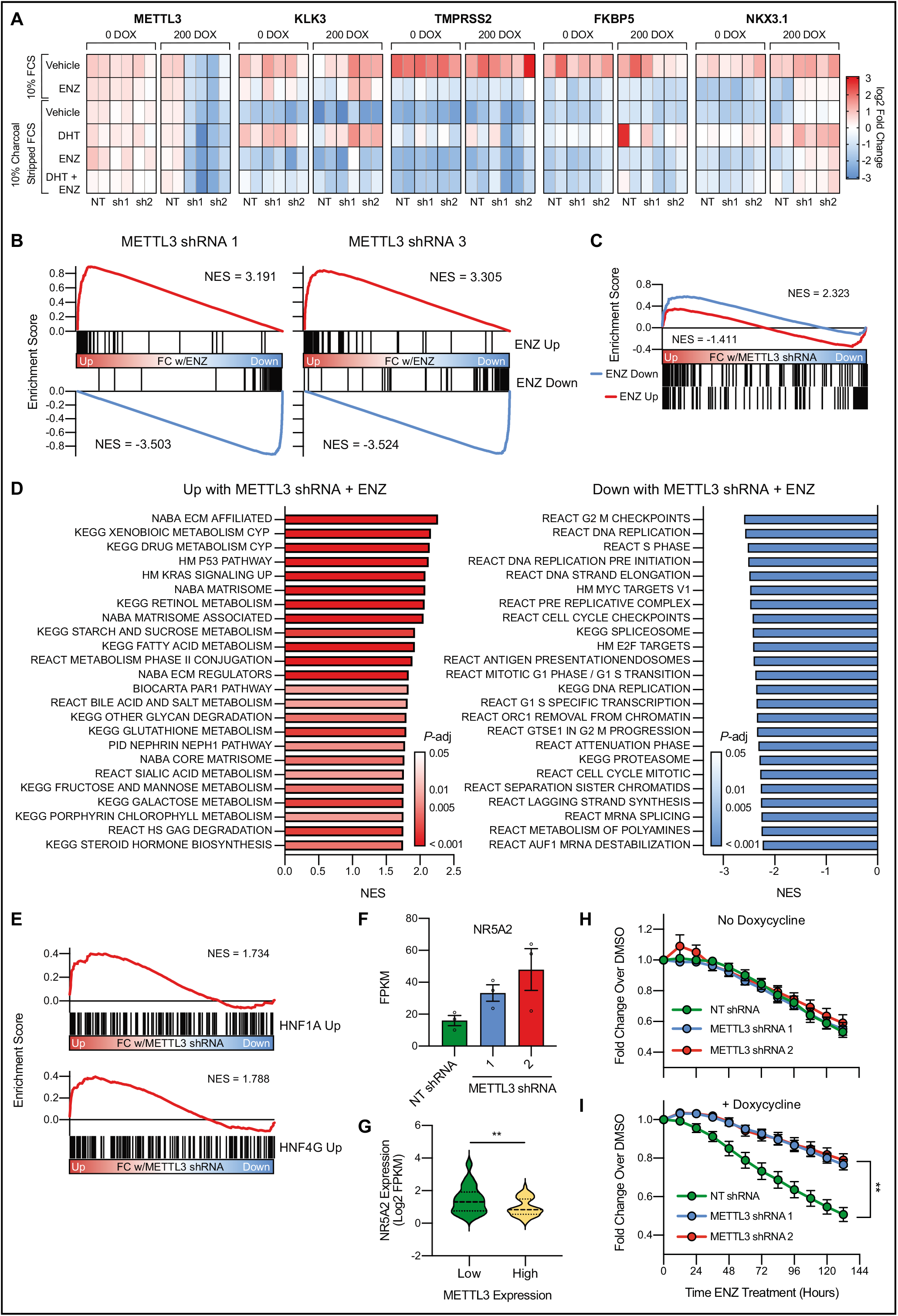
*METTL3* knockdown leads to the induction of a gene signature linked to castration resistance and renders the cells resistant to enzalutamide in an androgen receptor independent manner. **A)** Change in expression of AR target genes with *METTL3* knockdown (+/- doxycycline) and in response to AR stimulation (10 nM DHT) or inhibition (10 µM ENZ). Expression was measured via qPCR (n = 2) and is displayed as the fold change with treatment per gene and experiment. **B)** RNA-seq analysis of *METTL3* knockdown lines treated with ENZ demonstrates no change in the repression or activation of known ENZ target genes by gene set enrichment analysis. Genes were ranked according to the fold change ENZ vs vehicle as determined by DESeq2. ENZ up and down gene lists contain the common differentially expressed genes from two published RNA-seq experiments (GSE110903 and GSE147250). **C)** Some ENZ responsive genes show differential expression with *METTL3* knockdown. Ranking for gene set enrichment analysis was according to the combined fold change ENZ vs vehicle as determined by DESeq2 for both *METTL3* shRNA lines. **D)** Gene set enrichment analysis of genes differentially expressed with ENZ treatment and *METTL3* knockdown. Genes were ranked according to the combined Wald statistic from DESeq2 for both *METTL3* shRNA lines as compared to the NT shRNA. Shown are the top 24 results in either direction. **E)** *METTL3* knockdown with ENZ treatment induces the expression of HNF1A and HNF4G target genes. Genes were ranked according to the combined Wald statistic from DESeq2 for both *METTL3* shRNA lines as compared to the NT shRNA. HNF target genes were taken from previously published overexpression experiments in LNCaP cells (GSE85559). **F)** NR5A2 is overexpressed with *METTL3* knockdown and ENZ treatment (n = 3, error bars = SEM) **G)** NR5A2 expression in *METTL3* high vs. low samples (see Figure 1). Median and quartiles are indicated with a dotted line, *P* = 0.0034 as determined by t-test. **H)** In the absence of *METTL3* knockdown the shRNA lines respond similarly to 10 µM ENZ treatment (n = 3, error bars = SEM). **I)** Induction of *METTL3* knockdown renders the cells resistant to ENZ treatment (n = 4, error bars = SEM; sh1 *P* = 0.0012, sh2 *P* = 0.0011, as determined by t-test). For both panels shown is the change in cell confluency as compared to DMSO with time as measured via Incucyte imaging.

In order to get a global view of the response to *METTL3* knockdown and AR inhibition with ENZ, we analyzed changes in gene expression with RNA-seq. As expected, we again confirmed that *METTL3* knockdown did not change known AR signaling pathways, as most canonical ENZ responsive genes responded as expected to the treatment (Figure 5B). Nonetheless we did identify some genes, which similar to *NKX3*.*1*, demonstrated a reduction in the magnitude of their repression or activation with ENZ (Figure 5C). For example, *MAML2* is down-regulated 5.8-fold with ENZ and the NT shRNA, yet only 3.48-and 2.92-fold with the *METTL3* shRNAs (Supp Figure 5).

Overall 54 genes were significantly up-regulated with both *METTL3* shRNAs and under ENZ treatment conditions, while 25 genes were significantly down-regulated (FDR < 0.05, Supplementary Data 4). Importantly the upregulated genes included many which have been previously linked to ENZ resistance including: *TMEFF2, ADAMTS1*^47^, and *COL5A2*^48^. Gene set enrichment analysis of genes which are differentially expressed with *METTL3* knockdown in response to ENZ again identified an enrichment in pathways related extracellular matrix components (Figure 5D), which is consistent with our above results in patient expression data (Figure 1D), m^6^A mapping (Figure 2C), ribosome footprinting (Figure 3F), and proteomics (Figure 4D) analysis.

Additionally, we also identified many pathways involved in the metabolism of drugs and other compounds that were significantly upregulated with *METTL3* knockdown, again consistent with expression results from metastatic patient samples (Figure 1D). These same pathways, in addition to pathways that include complement and coagulation factors (which we see upregulated with *METTL3* knockdown Figure 1D/4D), make up a gastrointestinal-lineage transcriptional signature previously reported in PCa by Shukala, et al. This signature is regulated by HNF1A and HNF4G, and importantly leads to castration resistance^49^. Upon closer examination of our RNA-seq data, we specifically see a significant upregulation in genes which are upregulated by both HNF1A and HNF4G in LNCaP cells^49^ (Figure 5E). Although we do not see changes in the expression of either of these two transcription factors, we do see an upregulation of *NR5A2* (Figure 5F), an important gastrointestinal transcription factor which has been shown to regulate the same pathways in breast and colon cancer cells^50,51^. Examination of the patient data presented in Figure 1, also revealed that there is an inverse correlation with *NR5A2* and *METTL3* expression in CRPC patient samples (Figure 5G).

Given these results we then chose to examine how the *METTL3* knockdown lines respond at the cellular level to ENZ treatment. In the absence of *METTL3* knockdown all three cell lines responded similarly to 10 µM ENZ treatment, demonstrating an approximately 50% reduction in cell proliferation as compared to DMSO controls after 132 hours (Figure 5H). Strikingly, upon doxycycline-induced expression of the *METTL3* shRNAs, both lines showed a significant (sh1, *P* = 0.0012; sh2, *P* = 0.0011) resistance to the ENZ treatment with only an 80% reduction in proliferation after 132 hours (Figure 5I). Taken together these findings suggest that knockdown of *METTL3* promotes ENZ resistance through the upregulation of a gene signature driven by *NR5A2*.

## DISCUSSION

Current understanding of the effects of m^6^A is that the mark mostly serves to destabilize mRNAs^16,45^. As such, we would expect to see increases in the expression of methylated genes with *METTL3* knockdown. Strikingly, in our data we see relatively few changes in mRNA expression with *METTL3* knockdown. Furthermore, while changes in the translation of methylated transcripts are generally rarer, they predominantly serve to increase TE. Again, we would then expect to see a decrease in TE, whereas we see many increases at the level of TE and protein with *METTL3* knockdown in PCa. It therefore remains likely that we are also identifying some secondary-level changes due to alterations in the expression of upstream m^6^A methylated genes. It should also be noted that overall, our expression changes are fewer and more modest than many published studies. We propose that this is likely due to the fact that we are engineering an ∼70-80% knockdown instead of a complete *METTL3* knockout. While it may be possible to see stronger effects on gene expression with a *METTL3* knockout this phenotype is not clinically relevant as there is no complete loss of *METTL3* seen in PCa patients.

When examining pathways which are regulated by m^6^A in PCa, we consistently saw enrichment for extracellular matrix (ECM) proteins in both our cell line models (Figure 3F/4D/5D) and in CRPC patients (Figure 1D) with low levels of *METTL3*. Changes in the ECM are associated with increased tumor growth, migration and invasion, allowing for metastatic progression^52,53^, including in PCa^54^. Furthermore, changes in the ECM have been implicated in therapy resistance in breast, ovarian and PCa^55-58^. As such, it remains possible that these changes also contribute to the ENZ resistance demonstrated by the *METTL3* knockdown cells, and this would be a focus of future studies, in particular in a 3-D cell culture environment.

Lastly, our analysis of CRPC patient data demonstrated an inverse correlation between the levels of *METTL3* and AR signaling. When we examined the interplay between m^6^A and AR signaling we showed that though AR target genes and AR itself have relatively high numbers of m^6^A sites knockdown of *METTL3* did not change expression of these genes at the transcript or protein level. This may be due to low stoichiometry of the m^6^A sites on these transcripts (unresolved by miCLIP technology), redundancy with other components of the methyltransferase complex, or a lack of functionality of m^6^A on these transcripts. Nonetheless, we did find that knockdown of *METTL3* led to an upregulation of a previously identified gastrointestinal-lineage signature and rendered the cells resistant to ENZ in an AR-independent manner. Previously, this signature was shown to be driven by HNF4G and HNF1A which were induced in response to androgen deprivation^49^. Here we hypothesize that this signature is driven by an upregulation of another GI-lineage transcription factor *NR5A2* in response to *METTL3* knockdown. Although *NR5A2* itself was not identified as a methylated transcript in our miCLIP analysis, it is worth noting that the basal expression of *NR5A2* in LNCaP cells is relatively low. Therefore, it is possible that any potential methylation of this transcript may evade detection with an IP-based approach. Nonetheless it also plausible that *METTL3* knockdown is altering the methylation a yet unidentified upstream regulator of *NR5A2* as opposed to the gene itself. The exact mechanism behind how this signature leads to ENZ resistance remains to be determined. However, among the included genes are those involved in both the metabolism of steroid hormones (i.e. *AKR1C3* and *UGT2B15*) or general drug metabolism (i.e. *GSTA1, GSTA2, GSTK1*) both of which would play a probable role in ENZ resistance.

In summary, in this study we identified many genes and pathways which are dysregulated in the context of low *METTL3* expression in PCa. In particular we have nominated extracellular matrix proteins as being highly influenced by changes in *METTL3* expression. Furthermore, we showed that combining *METTL3* knockdown and ENZ treatment led to the upregulation of a GI-specific gene signature and rendered the cells resistant to ENZ. Future work will focus on delineating the precise mechanism behind the ENZ resistant phenotype in an attempt to determine whether *NR5A2* and/or other downstream pathway genes may function as potential therapeutic targets in CRPC. Overall, these findings support a new role for m^6^A in regulating therapeutic sensitivity to ARSi, and furthermore suggests that patients with low levels of *METTL3* expression may differentially respond to ARSi.

## METHODS

### Cell Lines

LNCaP cells (male, ATCC, RRID: CVCL_1379) were maintained in RPMI medium (Gibco, A1049101), supplemented with 10% FBS (Gibco, 10270106), and 1% penicillin-streptomycin (Gibco, 11548876) on poly-L-lysine coated plates. RWPE cells (male, ATCC, RRID: CVCL_3791) were maintained in Keratinocyte Serum Free Medium (Gibco, 17005075) supplemented with bovine pituitary extract and human recombinant EGF (included), and 1% penicillin-streptomycin (Gibco, 11548876). HEK293T cells (female, ATCC, RRID: CVCL_0063) and DU145 cells (male, ATCC, RRID: CVCL_0105) were maintained in DMEM (Gibco, 31966021), supplemented with 10% FBS, and 1% penicillin-streptomycin. All cell lines were grown at 37 °C with 5% CO_2_. All cell lines were authenticated by STR analysis and regularly tested for mycoplasma.

### m^6^A individual-nucleotide-resolution cross-linking and immunoprecipitation (miCLIP)

Total RNA from LNCaP and RWPE cells was isolated with TRIzol according to the manufacturer instructions. PolyA+ mRNA was isolated using the PolyATtract mRNA Isolation System (Promega, Z5210). 20 µg of mRNA was used as input for miCLIP following the previously reported protocol ^33^ and an m^6^A antibody from Abcam (ab151230). In parallel total mRNA was subjected to library preparation using the Illumina library preparation protocol. Final libraries were sequencing on an Illumina HiSeq2500 generating 50 bp paired-end reads at the Weill Cornell Medicine Epigenetic Core facility. In parallel total mRNA was subjected to library preparation using the Illumina library preparation protocol.

### Generation of inducible knock-down lines

Tet-pLKO-puro was a gift from Dmitri Wiederschain (Addgene plasmid # 21915; http://n2t.net/addgene:21915 ; RRID: Addgene_21915). Tet-pLKO-puro was digested with AgeI and EcoRI and ligated with annealed oligos (Supplementary Table 1). Lentivirus was produced in HEK293T cells by transfection with the pLKO constructs, and subsequent virus containing media was used to transduce LNCaP cells. Three days post transduction the cells were subjected to puromycin selection (1 µg/mL).

### RNA-and Ribo-Seq

For paired RNA and Ribosome profiling experiments cells (∼15×10^6^) were incubated with 100 µg/ml cycloheximide for 5 minutes at 37□°C. Cells were then trypsinized, pelleted, and the pellets washed twice with ice-cold PBS containing 100 µg/ml cycloheximide. An aliquot was set aside for confirmation of knock-down by western blot, and the remaining cells were resuspended in 425 µl of hypotonic buffer (5 mM Tris-HCl (pH 7.5), 2.5 mM MgCl2, 1.5 mM KCl and 1x Halt protease inhibitor cocktail (Thermo Scientific, 78410)), followed by the addition of 50 µg cycloheximide, 1 µl of 1M DTT, and 100 units of RNAse Inhibitor (Thermo Fisher Scientific, N8080119). The pellet was vortexed, 25 µl 10% Triton X-100 and 25 µl 10% sodium deoxycholate were added then followed by another vortex. The lysates were then cleared by spinning at 16,000 g for 8 minutes at 4□°C. The lysate was diluted 1:10 and OD at 260 nm was determined using a Nanodrop spectrophotometer.

Ribosome footprinting was based on the manufacturer’s instructions for the TruSeq Ribo Profile (Mammalian) Library Prep Kit (Illumina, discontinued). Briefly, lysates were normalized to OD of 30 and a volume of 200 µl. Then 3 µl of RNAse I (Ambion, AM2294) was added and the samples were incubated for 45 minutes at room temperature with shaking. 10 µl of SUPERase•In was added to stop the reaction. Ribosome bound RNA fragments were purified on MicroSpin S-400 HR Columns (Cytiva, 27-5140-01) followed by cleanup with the RNA Clean & Concentrator-25 kit. rRNA was removed using the NEBNext rRNA Depletion Kit (NEB, E6310), and 28-30 bp footprints were purified on a 15% polyacrylamide TBE/Urea gel. Ribo-seq libraries were then prepared with the purified footprints and the SMARTer Small RNA-Seq Kit (Takara, 635029) following the manufacturer’s instructions.

RNA was extracted from either 100 µl of ribosome footprinting lysate with the RNA Clean & Concentrator-25 kit (Zymo, R1017), or directly from cells using the RNeasy Mini Kit (Qiagen, 74106), and genomic DNA was removed using the DNA-free kit (Ambion, AM1906). RNA quality was assessed with a BioAnalyzer and quantity on a Qubit fluorometer. RNA-seq libraries were prepared using 1 µg of RNA and the NEBNext Ultra^™^ II RNA Library Prep Kit (NEB, E7775) with rRNA depletion following the manufacturer’s instructions.

All libraries were sequenced on an Illumina NextSeq generating 75 bp single-end reads at University of Basel Visceral Surgery and Precision Medicine Research Laboratory.

### Bioinformatics Analysis of Sequencing Data

#### miCLIP

Crosslinking-induced mutation sites were identified in miCLIP datasets as described previously ^33^. Individual m^6^A sites were subjected to metagene analysis using MetaPlotR ^59^. These identified sites were further filtered only reporting those sites that had greater than one crosslinking-induced mutation (m ≥ 2) and those that mapped to an A residue to generate Supplementary Data 1.

#### RNA-Sequencing

Sequence reads were aligned to the human reference genome GRCh37 by STAR using the two-pass approach ^60^. Transcript quantification was performed using RSEM ^61^. Genes without >10 counts in at least 2 samples were discarded. Counts were normalized using the median of ratios method from the DESeq2 package in R version 3.6.1 (R Core Team (2019). R: A language and environment for statistical computing. R Foundation for Statistical Computing, Vienna, Austria. URL https://www.R-project.org/). Differential expression analysis was performed using the wald test in DESeq2 ^62^.

#### Ribo-Sequencing

Sequence reads were aligned as for RNA-seq, using STAR, but with the options – “winAnchorMultimapNmax 200 −-seedSearchStartLmax 15 --outFilterMultimapNmax 20”, to improve mapping of short Ribo-seq reads. Differential analysis of ribosome loading was performed using Riborex which uses generalised linear models and the DESeq2 normalisation framework to identify genes showing altered translation between sample groups ^40^.

### Drug Treatments

For all drug treatments cells were pre-treated with 200 ng/ml doxycycline for 96 hours to induce shRNAs. For DHT stimulation experiments cells were starved of hormone for 48 hours in phenol red-free RPMI media (Gibco, 11-835-030) with 10% charcoal stripped FBS (Gibco, A3382101), then treated with 10 nM dihydrotestosterone or 10 µM enzalutamide for 24 hours. For Incucyte experiments cells were treated with doxycycline for 96 hours and then plated in a 96-well plate, 5000 cells per well, n = 8 per condition. Cell confluency was determined using the Incucyte S3 instrument and the IncuCyte S3 2018B software. Values were calculated as fold-change in confluency as compared to vehicle treated controls.

### Proteomics

Inducible shRNA lines were treated with 200 ng/mL doxycycline for 96 hours. Cells were scraped directly in lysis buffer (8M urea, 100mM Tris pH8, plus protease inhibitors) and lysed with sonication for 1 minute on ice with 10 seconds intervals. The supernatant was reduced, alkylated and precipitated overnight. The pellet was re-suspended in 8M urea/50mM Tris pH8 and protein concentration was determinate with Qubit Protein Assay (Invitrogen).

10 µg of protein were digested with LysC for 2 hours at 37□°C followed by Trypsin at room temperature overnight. 800 ng of digests were loaded in random order onto a pre-column (C18 PepMap 100, 5 µm, 100 A, 300 µm i.d. x 5 mm length, Thermo Fisher, 160454) at a flow rate of 50 µL/min with solvent C (0.05% TFA in water/acetonitrile 98:2).

After loading, peptides were eluted in back flush mode onto a home packed analytical Nano-column (Reprosil Pur C18-AQ, 1.9 µm, 120 A, 0.075 mm i.d. x 500 mm length) using an acetonitrile gradient of 5% to 40% solvent B (0.1% Formic Acid in water/acetonitrile 4,9:95) in 180 min at a flow rate of 250 nL/min. The column effluent was directly coupled to a Fusion LUMOS mass spectrometer (Thermo Fischer, Bremen; Germany) via a nano-spray ESI source.

Data acquisition was made in data dependent mode with precursor ion scans recorded in the orbitrap with resolution of 120’000 (at m/z=250) parallel to top speed fragment spectra of the most intense precursor ions in the Linear trap for a cycle time of 3 seconds maximum.

The mass spectrometry data were processed with MaxQuant ^63^ (version 1.6.1.0) against the Swissprot Homo Sapiens database ^64^ (release February 2019). The initial precursor mass tolerance was set to 10 ppm, and that of the fragment peaks to 0.4 Da. Enzyme specificity was set to strict trypsin, and a maximum of three missed cleavages were allowed. Carbamidomethylation on cysteine was set as a fixed modification, methionine oxidation and protein N-terminal acetylation as variable modifications. The same fraction number was given to all replicates of a group, but each group was given a fraction number differing by 3, so that match between runs were prevented between the different groups.

Protein intensities are reported as MaxQuant’s Label Free Quantification (LFQ) values, as well as iTop3 ^65^ values (sum of the intensities of the three most intense peptides); for the latter, variance stabilization ^66^ was used for the peptide normalization, and missing peptide intensities, if at least 2 evidences exist in a group, were imputed by drawing values from a Gaussian distribution of width 0.3 centered at the sample distribution mean minus 1.8x the sample standard deviation. Imputation at protein level for both iTop3 and LFQ was performed if there were at least two measured intensities in at least one group of replicates; missing values in this case were drawn from a Gaussian distribution of width 0.3 centered at the sample distribution mean minus 2.5x the sample standard deviation. Differential expression tests were performed using empirical Bayes (moderated t-test) implemented in the R limma package ^67^. The Benjamini and Hochberg ^68^ method was further applied to correct for multiple testing.

### Immumoblotting

Cells were lysed in RIPA buffer with protease inhibitors, resolved on 4-15% Mini-Protean TGX gels (BioRad, 456-1084), and transferred to nitrocellulose membranes using the iBlot2 system (Thermo Fisher, IB23001). Blots were blocked for 1 hour at room temperature and then probed with primary antibodies against METTL3 (Cell Signaling, 96391), KIF5C (Abcam, ab193352), GAPDH (Millipore, AB2302), IFIT3 (Santa Cruz, sc-393512), STAT1 (Cell Signaling, 14994), phospho-STAT1 (Cell Signaling, 7649), or beta-actin (Abcam, ab6276) overnight in 5% milk or BSA (all Cell Signaling antibodies) in TBST at 4□°C. After 4 washes, blots were incubated with IRDye secondary antibodies (LI-COR, 926-68075, 926-32211, 926-32210) in 5% milk for one hour at room temperature. After 4 washes blots were visualized on a LI-COR Odyssey Infrared Imaging System.

### qPCR

RNA was extracted directly from cells using the RNeasy Mini Kit with an on column DNAse treatment. RNA was reverse transcribed using the FIREScript RT cDNA kit and random primers (Solis Biodyne, 06-15-00200) according to the manufacturer’s instructions. Quantitative real-time PCR was performed on the ViiA 7 system (Applied Biosystems) using HOT FIREPol EvaGreen qPCR mix (Solis Biodyne, 08-24-00020) following manufacturer’s instruction. Primer sequences are listed in Supplementary Table 1. All quantitative real-time PCR assays were carried out using three technical replicates. Relative quantification of quantitative real-time PCR data used GAPDH, actin, and HMBS as housekeeping genes.

### Interferon Stimulation

Cells were treated with 200 units/mL of recombinant interferon alpha (Abcam, ab48750) for between 4 and 24 hours before Western blot analysis.

## RESOURCE AVAILABILITY

### Lead Contact

Further information and requests for resources and reagents should be directed to and will be fulfilled by the Lead Contact, Mark Rubin.

### Materials Availability

Inducible shRNA plasmids generated in this study have been deposited to Addgene (Plasmid #: 162984, 162985, and 163017). Cell lines generated in this study are available from the Lead Contact with a completed Materials Transfer Agreement.

### Data and Code Availability

The miCLIP, RNA, and Ribo-Seq data discussed in this publication have been deposited in NCBI’s Gene Expression Omnibus ^69^ and are accessible through GEO Series accession number GSE161304 (https://www.ncbi.nlm.nih.gov/geo/query/acc.cgi?acc=GSE161304).

Raw images of blots in the paper have been deposited to Mendeley Data: http://dx.doi.org/10.17632/xv2vzc8pzz.1

The mass spectrometry proteomics data have been deposited to the ProteomeXchange Consortium (http://proteomecentral.proteomexchange.org) via the PRIDE partner repository^70^ with the dataset identifier PXD022348.

## ACKNOWLEDGEMENTS

We thank the members of the Rubin laboratory for their comments and suggestions. We thank the Weill Cornell Epigenomics Core and Viola Paradisio (Department of Biomedicine, University of Basel) for their assistance in running the high-throughput sequencing. We also thank the University of Bern Proteomics Core for their assistance with the proteomics experiments. We acknowledge expert assistance from Mariana Ricca at the University of Bern in preparing the manuscript for submission. This project was supported by funding from the Prostate Cancer Foundation (18YOUN06, K.A.C.), the Weill Cornell Medicine SPORE in Prostate Cancer (Developmental Research Project, S.R.J. and M.A.R.), the Swiss Cancer League (KFS-4988-02-2020-R, S.P.), and the NIH (R01CA186702, S.R.J.).

## AUTHOR CONTRIBUTIONS

K.A.C., M.A.R., and S.R.J. designed the study and the experiments. K.A.C., N.U., and P.R. performed the experiments. K.M. performed the miCLIP library prep and data analysis. J.G. and S.P. performed the data analysis for the RNA-seq and ribosome footprinting. K.A.C., M.A.R., and S.R.J. wrote the initial draft. All authors reviewed and edited the manuscript.

