## Supplemental Material for "Mapping of m^6^A and Its Regulatory Targets in Prostate Cancer Reveals a METTL3-low Induction of Therapy Resistance"

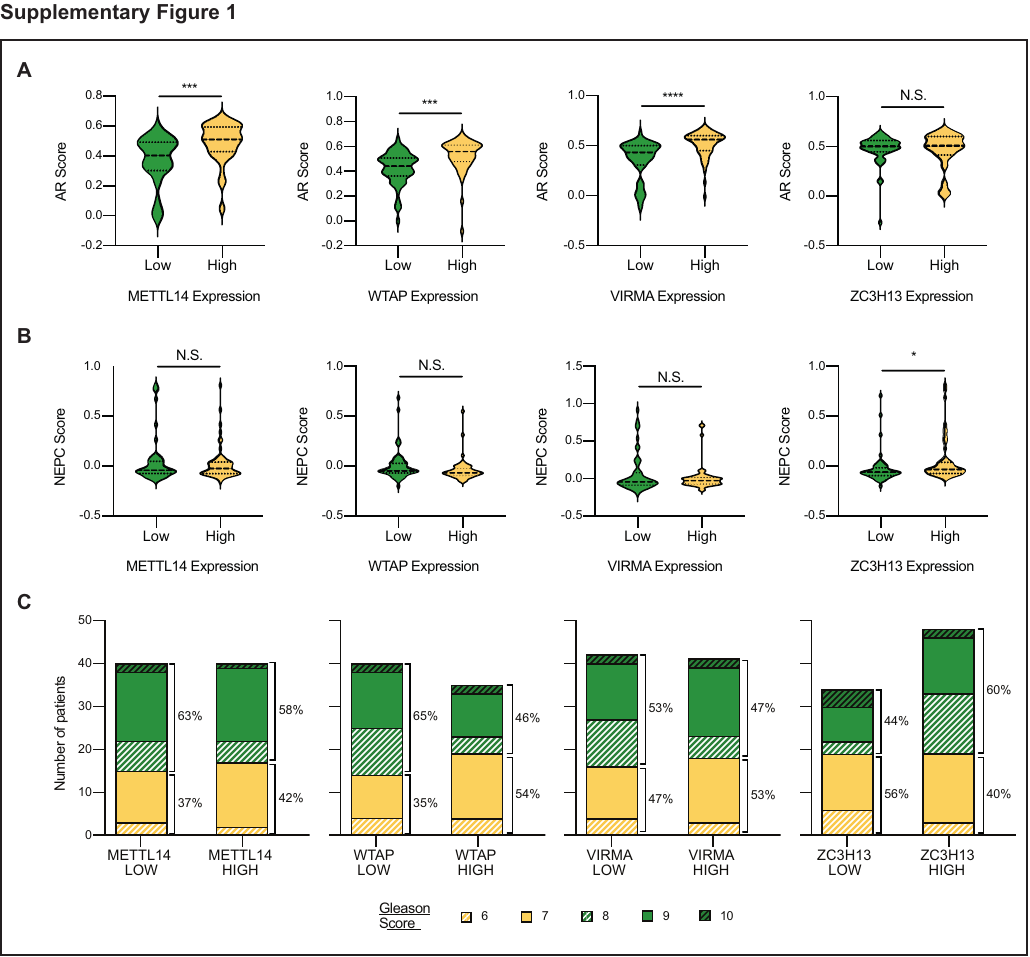

**Supplementary Figure 1. Association between m^6^A “writer” proteins and markers of advanced-stage disease. A)** Castration resistant prostate cancer samples^31^ were grouped based on *METTL14*, *WTAP*, *VIRMA*, or *ZC3H13* expression: high, z-score > 1, n = 46, 50, 52, and 48 respectively; low, z-score < -1, n = 57, 50, 55, and 52 respectively. **A)** AR score in high vs. low samples. Median and quartiles are indicated with a dotted line, statistical significance is indicated as determined by t-test. **B)** NEPC score in in high vs. low samples. Median and quartiles are indicated with a dotted line, statistical significance is indicated as determined by t-test. **C)** Gleason score in high vs. low samples. No statistically significant differences were detected by Fischer’s exact test between samples with Gleason 6-7 and Gleason 8+ scores.

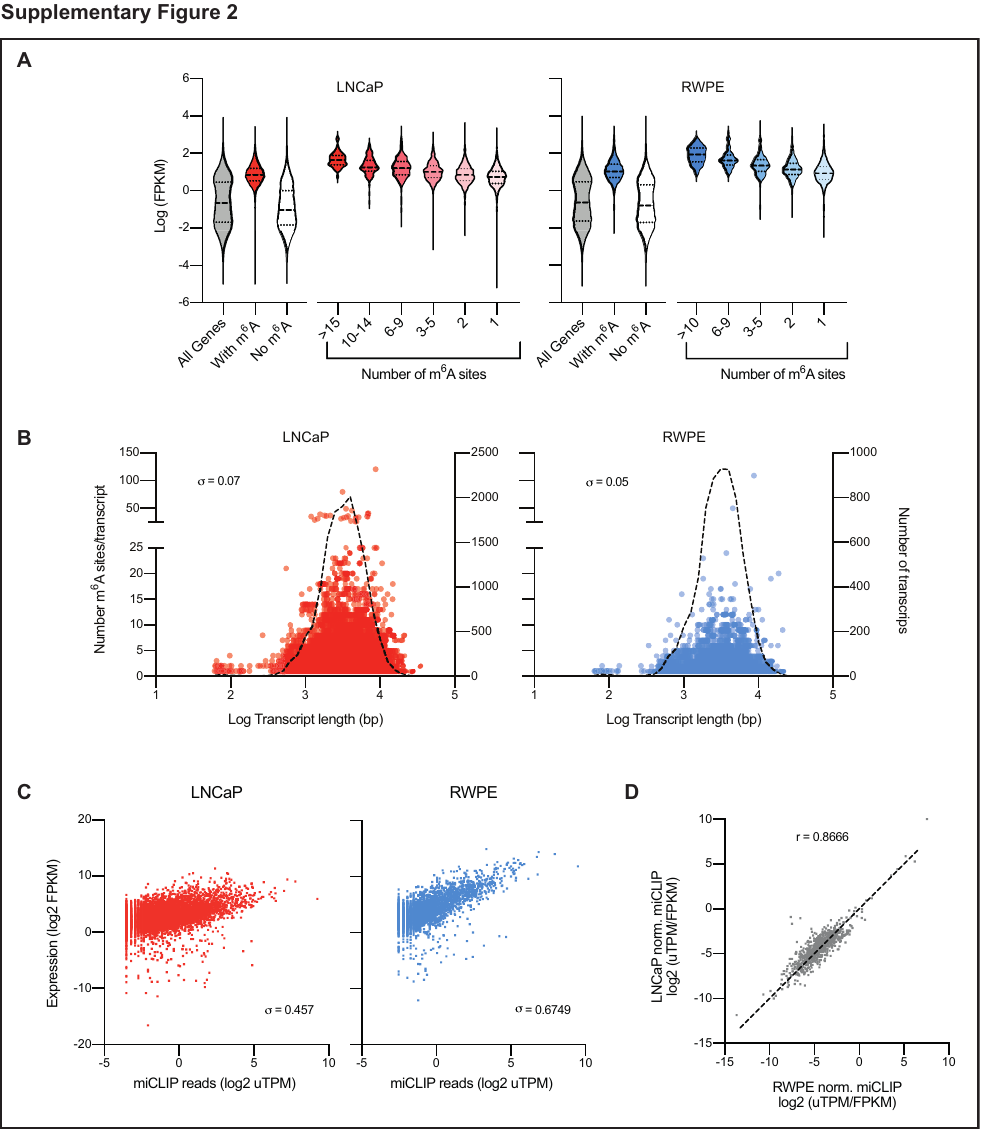

**Supplementary Figure 2. Relation of miCLIP results to transcript length and expression. A)** Comparing the number of m^6^A sites per transcript to the transcript expression as determined by paired RNA-Seq. **B)** Comparing the number of m^6^A sites per transcript (circles, plotted on the left y-axis) to the transcript length shows no correlation (Spearman). The distribution is similar to the overall number of transcripts of a given length (dotted line, plotted on the right y-axis). **C)** Normalizing m^6^A levels per transcript (unique tags per million, uTPM) by the transcript expression from RNA-seq (FPKM). **D)** Correlation between the normalized m^6^A levels per site (uTPM/FPKM) for the sites common to both LNCaP and RWPE cells lines (n = 889) (Pearson r = 0.937).

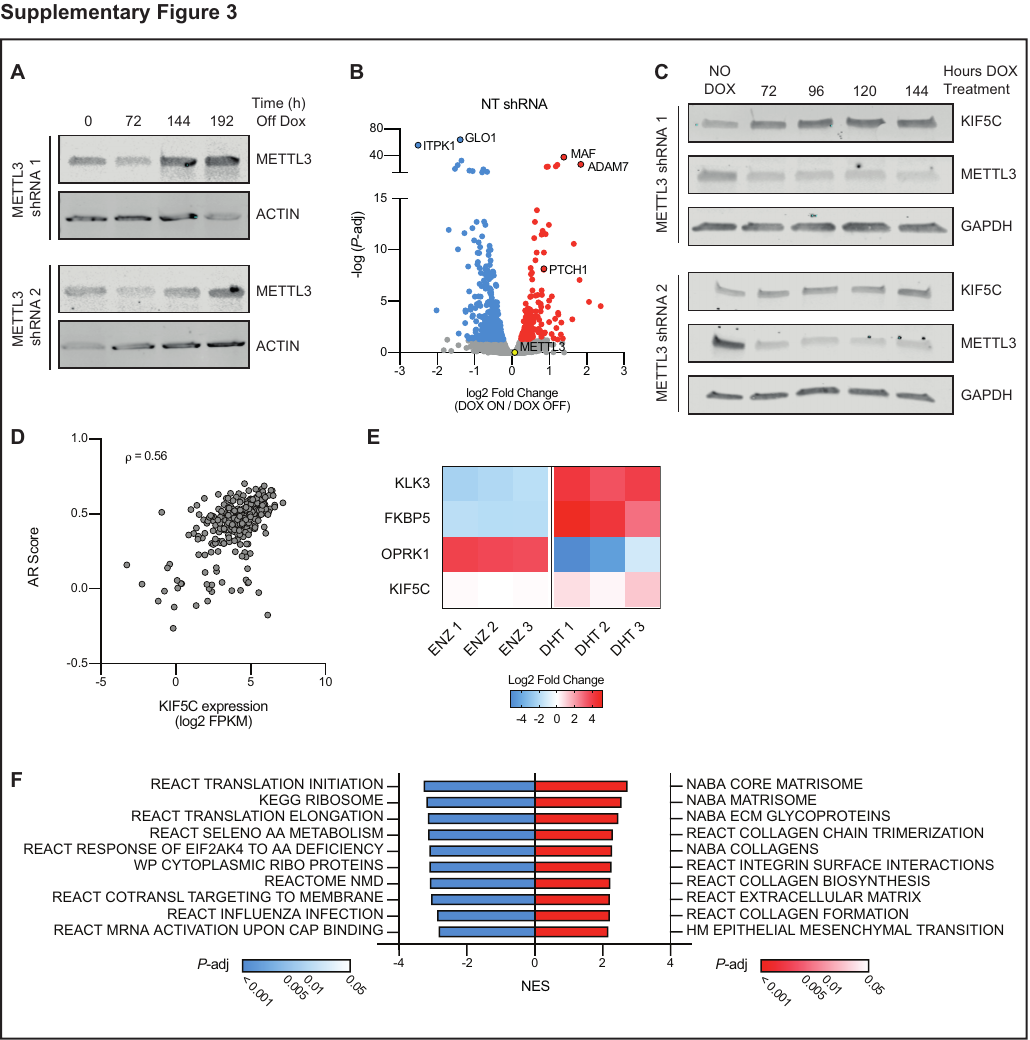

**Supplementary Figure 3. Reversible knockdown of *METTL3* in LNCaP cells upregulates KIF5C. A)** Doxycycline induction of *METTL3* shRNA is reversible. Cells were treated with doxycycline for 96 hours, and then changed to media without doxycycline for the indicated amount of time followed by Western blot analysis for METTL3. **B)** Significantly (adjusted *P*-value < 0.05) differentially expressed genes with induction of the NT shRNA (n = 3). Common genes also induced in the two *METTL3* shRNA lines are highlighted with a black border. **C)** Western blot analysis of the upregulation of KIF5C with METTL3 knockdown. Cells were treated with doxycycline for the indicated amount of time. **D)** *KIF5C* expression is correlated with AR score in castration resistant prostate cancer samples^31^ (n = 264) (Spearman’s ρ = 0.56). **E)** Fold change in expression of the *KIF5C* and the AR-regulated genes *KLK3, FKBP5* and *OPRK1* with DHT or ENZ treatment of LNCaP cells in published RNA-seq datasets (GSE110903, GSE147250, GSE135879, GSE130534, GSE115395).

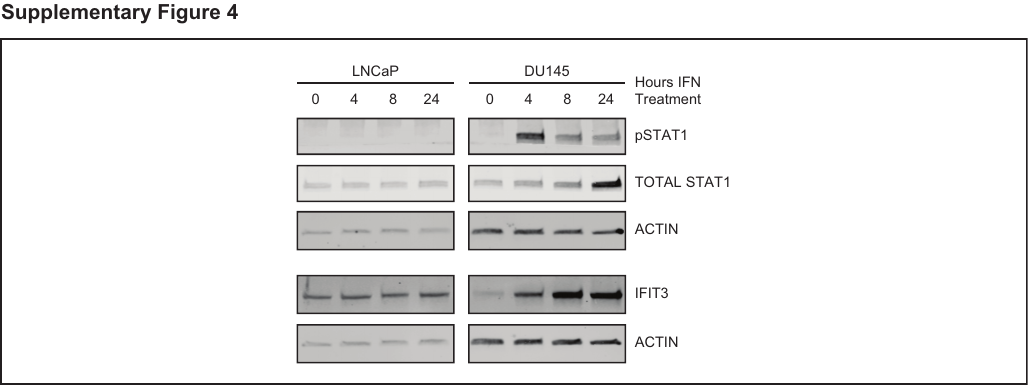

**Supplementary Figure 4. LNCaP cells do not respond to IFN.** Western blot analysis of IFN response as determined by the upregulation of STAT1 and IFIT3 in LNCaP and DU145 cells in response to IFN-α treatment.

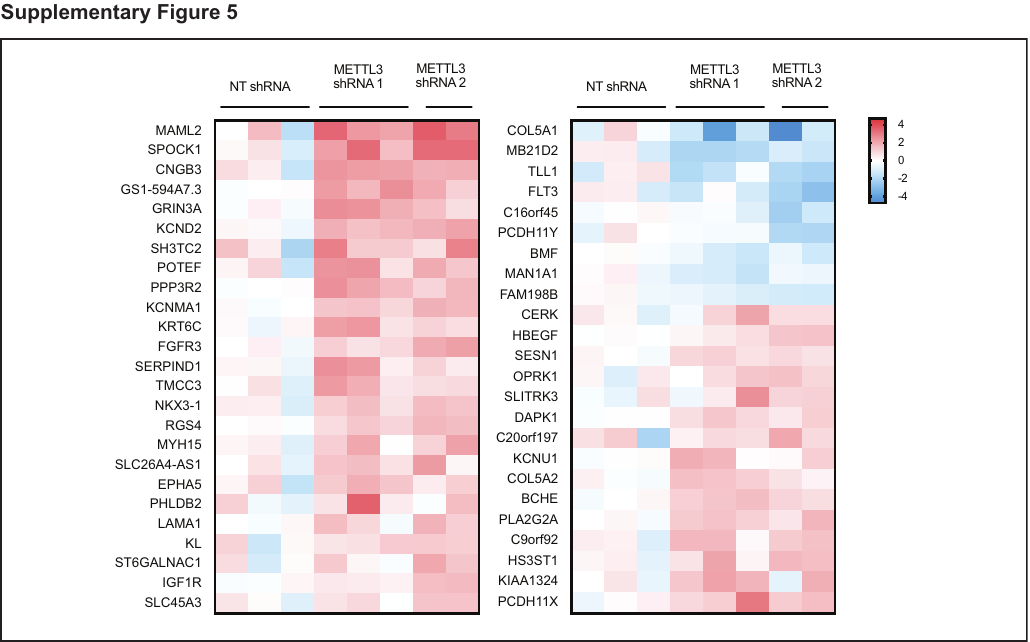

**Supplementary Figure 5. A subset of ENZ-regulated genes are differentially expressed with METTL3 knockdown.** Data is expressed as the fold-change over NT shRNA for all three lines with 24 hours of 10 µM ENZ treatment. On the left and right are genes down-and up-regulated by ENZ respectively.

**Supplementary Table 1. Oligonucleotide Sequences**

| qPCR | GAPDH | Fwd | GAC AGT CAG CCG CAT CTT CT |
| --- | --- | --- | --- |
| qPCR | GAPDH | Rev | TTA AAA GCA GCC CTG GTG AC |
| qPCR | HMBS | Fwd | AGC TTG CTC GCA TAC AGA CG |
| qPCR | HMBS | Rev | AGC TCC TTG GTA AAC AGG CTT |
| qPCR | ACTB | Fwd | TGA CGT GGA CAT CCG CAA AG |
| qPCR | ACTB | Rev | CTG GAA GGT GGA CAG CGA GG |
| qPCR | METTL3 | Fwd | TTG TCT CCA ACC TTC CGT AGT |
| qPCR | METTL3 | Rev | CCA GAT CAG AGA GGT GTA G |
| qPCR | TMPRSS2 | Fwd | GTC CCC ACT GTC TAC GAG GT |
| qPCR | TMPRSS2 | Rev | CAG ACG ACG GGG TTG GAA G |
| qPCR | KLK3 | Fwd | TGG GGA CCA CCT GCT ACG CC |
| qPCR | KLK3 | Rev | TCG GTG ATC AGA ATG ACC CAC GAG |
| qPCR | FKBP5 | Fwd | GCA ACA GTA GAA ATC CAC CTG |
| qPCR | FKBP5 | Rev | CTC CAG AGC TTT GTC AAT TCC |
| qPCR | NKX3.1 | Fwd | CCC ACA CTC AGG TGA TCG AG |
| qPCR | NKX3.1 | Rev | GAG CTG CTT TCG CTT AGT CTT |
| shRNA | METTL3 1 | Fwd | CCGGGCAAGTATGTTCACTATGAAACTCGAGTTTCATAGTGAACATACTTGCTTTTTG |
| shRNA | METTL3 1 | Rev | AATTCAAAAAGCAAGTATGTTCACTATGAAACTCGAGTTTCATAGTGAACATACTTGC |
| shRNA | METTL3 2 | Fwd | CCGGGCTGCACTTCAGACGAATTATCTCGAGATAATTCGTCTGAAGTGCAGCTTTTTG |
| shRNA | METTL3 2 | Rev | AATTCAAAAAGCTGCACTTCAGACGAATTATCTCGAGATAATTCGTCTGAAGTGCAGC |
| shRNA | GFP | Fwd | CCGGTACAACAGCCACAACGTCTATCTCGAGATAGACGTTGTGGCTGTTGTATTTTTG |
| shRNA | GFP | Rev | AATTCAAAAATACAACAGCCACAACGTCTATCTCGAGATAGACGTTGTGGCTGTTGTA |
